# Universal attenuators and their interactions with feedback loops in gene regulatory networks

**DOI:** 10.1101/074716

**Authors:** Dianbo Liu, Luca Albergante, Timothy J Newman

**Affiliations:** School of Life sciences, University of Dundee, Dow Street, Dundee, DD1 5EH, UK; The Broad Institute of Harvard and MIT, 415 Main Street, Cambridge, MA 02142, USA; Computer Science and Artificial Intelligence Lab, Massachusetts Institute of Technology, 32 Vassar St, Cambridge, MA 02139, USA; Institut Curie, 26 Rue d’Ulm, 75005 Paris, France

## Abstract

Using a combination of mathematical modelling, statistical simulation and large-scale data analysis we study the properties of linear regulatory chains (LRCs) within gene regulatory networks (GRNs). Our modelling indicates that downstream genes embedded within LRCs are highly insulated from the variation in expression of upstream genes, and thus LRCs act as attenuators. This observation implies a progressively weaker functionality of LRCs as their length increases. When analysing the preponderance of LRCs in the GRNs of *E. coli* K12 and several other organisms, we find that very long LRCs are essentially absent. In both *E. coli* and *M. tuberculosis* we find that four-gene LRCs are intimately linked to identical feedback loops that are involved in potentially chaotic stress response, indicating that the dynamics of these potentially destabilising motifs are strongly restrained under homeostatic conditions. The same relationship is observed in a human cancer cell line (K562), and we postulate that four-gene LRCs act as “universal attenuators”. These findings suggest a role for long LRCs in dampening variation in gene expression, thereby protecting cell identity, and in controlling dramatic shifts in cell-wide gene expression through inhibiting chaos-generating motifs.

**In brief:** We present a general principle that linear regulatory chains exponentially attenuate the range of expression in gene regulatory networks. The discovery of a universal interplay between linear regulatory chains and genetic feedback loops in microorganisms and a human cancer cell line is analysed and discussed.

**Highlights:** Within gene networks, linear regulatory chains act as exponentially strong attenuators of upstream variation

Because of their exponential behaviour, linear regulatory chains beyond a few genes provide no additional functionality and are rarely observed in gene networks across a range of different organisms

Novel interactions between four-gene linear regulatory chains and feedback loops were discovered in *E. coli, M. tuberculosis* and human cancer cells, suggesting a universal mechanism of control.

## Introduction

The behaviour of cells is controlled in large part by the coordinated activation and inhibition of thousands of genes. This coordination is achieved via a complex network of gene regulation that enables a cell to express the appropriate set of genes for a particular environment and/or phenotype. The primary mode of gene regulation is through a class of genes that encode proteins which bind to regulatory regions on the DNA. These transcription factors (TFs) activate or inhibit the expression of typically a large number of downstream target genes. Genome-wide studies of TF binding allow the construction of gene regulatory networks (GRNs) that summarize the global structure of genetic interactions; each node represents a gene and an arrow between two nodes denotes the regulation of a target gene by a TF-coding gene (which we will describe for brevity as a TF unless there is potential for confusion). The dynamics of transcriptional regulation are still not fully understood [1].

However, over relatively long time scales, transcriptional response is generally analogue, i.e. a stronger expression of a TF gene results in a higher nuclear concentration of the TF protein and thereby a stronger activation or inhibition of the target genes [2-6].

GRNs typically contain thousands of genes and are beyond simple intuitive interpretation and understanding. Therefore, computational and mathematical approaches must be employed to gain a better understanding of the structure and function of system-level genetic interaction. One widely used approach focuses on the study of small-scale network configurations, called motifs [4, 7], and on their functional pressures. This approach has been effective in uncovering the functionality of motifs often encountered across different networks, such as the feed-forward loop and the bi-fan. The combinatorial complexity of GRNs limits the applicability of this analysis to motifs comprising more than four nodes, and complimentary ways of analysing networks are important to better understand how larger-scale topology is associated with GRN function [4, 8, 9].

In this article, we use a methodology inspired by motif analysis to study the behaviour of a particular class of network configurations that we call linear regulatory chains (LRCs). Our approach exploits the theoretical power of mathematical and statistical analysis to determine the expected behaviour of LRCs and to derive predictions that we then test on biological datasets available in the literature to obtain a better understanding of the selection pressures acting on GRNs.

For the purpose of our mathematical analysis, we define LRCs as linear chains of one-way regulation in which each node interacts with at most one node downstream and one node upstream. A given interaction can be either inhibitory or activating. Each LRC starts at the top layer (no transcriptional input) and ends at the bottom layer (no transcriptional output^1^) of the respective GRN. We relax this definition when studying real GRN datasets, and define LRCs as linear chains of genes which form a causal chain of transcriptional interaction, without placing restrictions on the number of connections to any given node.

While our analysis is focussed here on transcriptional interactions, the generality of network modelling allows the application of our results to other contexts in molecular biology and beyond [5, 10, 11].

## Results

### Mathematical formulation of gene regulation

To investigate the effect of LRCs in gene expression we employed a minimal mathematical model of transcriptional regulation. The model describes a linear chain of regulation as portrayed in Figure 1A. For two adjacent genes in the chain, we assume that the rate of transcription of the target gene varies smoothly with the concentration of the TF of the corresponding upstream gene, according to a Hill-like function (see Materials and Methods). This function is characterised by four parameters and is able to describe both activation and inhibition (Figure 1B-C and Figure S. 1A). To minimize the complexity of our model, we assume that protein concentration can be used as a proxy for gene expression, i.e. that the rate of transcription and the protein concentration are proportional. We define the source TF as the TF at the top of the chain which is not itself under transcriptional regulation. Most of our analysis concerns the effective regulation by the source TF on downstream genes. Therefore, it is very important to distinguish the regulation of a downstream node due to its immediate upstream TF and the effective regulation of the same node due to the source TF. A useful feature of the Hill-like function that we use is its universality: if it describes each of the individual links in the chain (with Hill coefficient of unity), then the net regulation of a node due to the source TF can also be described by the very same function, thus providing a simplified description of the LRC.

The effective regulation of the *n*^th^ gene in an LRC can be described by a function of the expression level of the source TF. This function will be called R_n_(-). The regulatory effect of R_n_(-) can be summarised by three quantities: the “lowest level of expression” (LLE), the “highest level of expression” (HLE), and the “effectiveness”, which is the difference between HLE and LLE. With regard to the last of these three, it is more convenient in our analysis to define “relative effectiveness” (RE), which is the difference between HLE and LLE divided by their mean (see Table 1 for a summary of the quantities introduced). More formally:

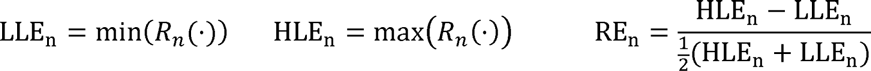

**Table 1.** Interpretation of the quantities introduced to describe transcriptional activation and inhibition

|  | Net transcriptional activation | Net transcriptional inhibition |
| --- | --- | --- |
| Lowest level of Expression (LLE) | Baseline transcriptional level, i.e. level of transcription in the absence of the activator. | Limit of transcriptional inhibition, i.e. minimum level of transcription when the concentration of the inhibitor is arbitrarily large. |
| Highest Level of Expression (HLE) | Limit of transcriptional activation, i.e. maximum level of transcription when the concentration of the activator is arbitrarily large. | Baseline transcriptional level, i.e. level of transcription in the absence of the inhibitor. |
| Relative Effectiveness (RE) | Maximum range of variation in the concentration of the target gene (Effectiveness) normalised by the mean of the extreme points. | Maximum range of variation in the concentration of the target gene (Effectiveness) normalised by the mean of the extreme points. |

These three measures complement the Hill coefficient, which is commonly defined for transcriptional response and determines the degree of non-linearity, i.e. “sensitivity”, of the regulatory function [12].

The precise sequence of inhibition and activation within an LRC dictates the net effect on a given target gene when the expression of the source TF is varied. For example, consider an LRC comprising only inhibitory interactions: after an even number of regulatory steps, the initial gene acts as an activator, while after an odd number it acts as an inhibitor (Figure 1D). On the other hand, when the LRC comprises only activating interactions, all downstream genes are effectively activated by the source TF. If the LRC is a mixture of activating and inhibiting interactions, the type of net regulation of a given gene depends on the number of upstream inhibitory links. Due to this dynamic diversity, we will focus primarily on LRCs comprising only inhibitory steps. The relevance of our analysis to more heterogeneous LRCs will be discussed as appropriate.

**Figure 1.**
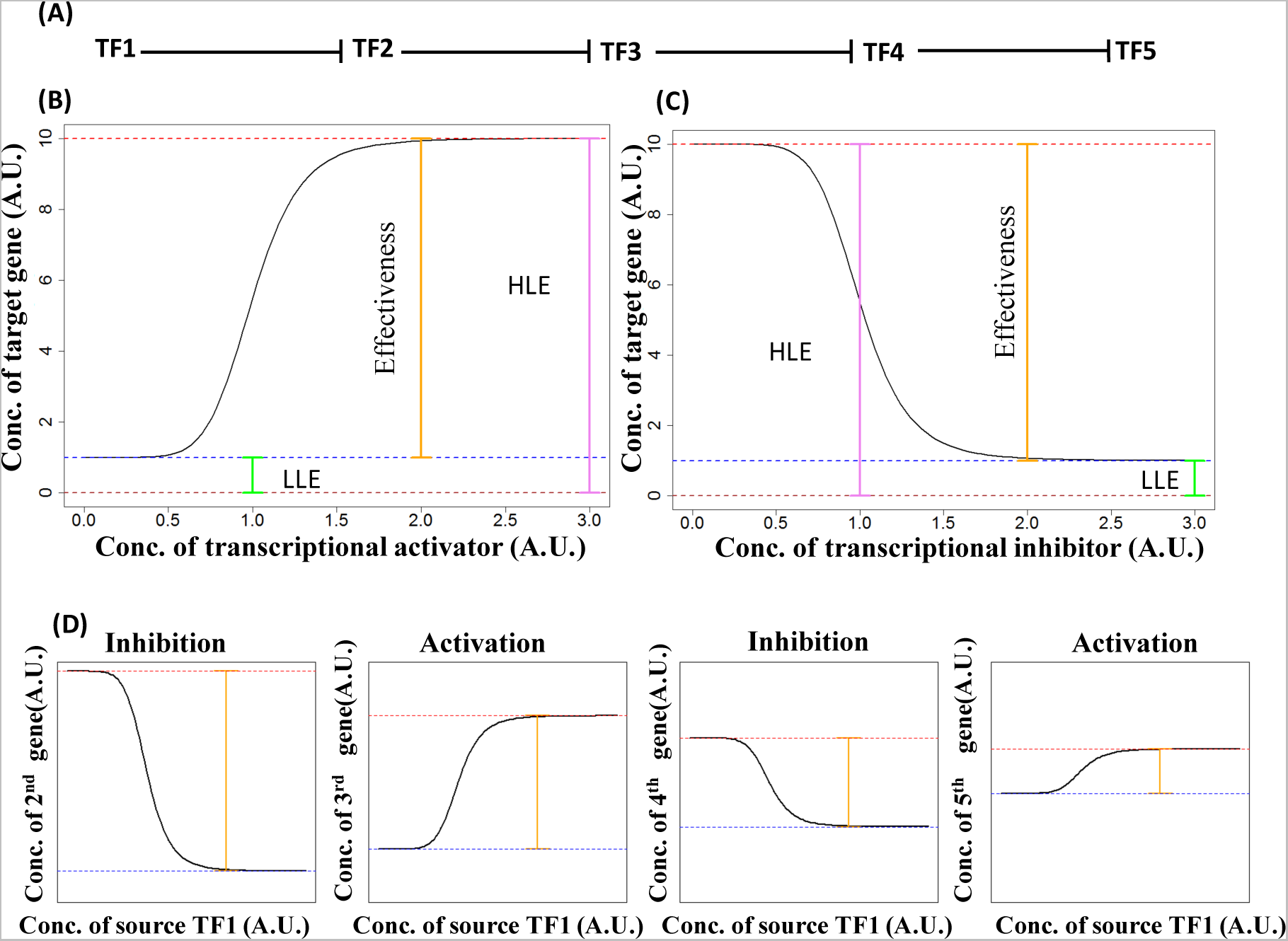
Dynamics of long regulatory chains (LRC). (A) Schematic of a 5 gene LRC formed by inhibitory transcriptional regulations. (B) Net transcriptional activation. The figures summarize the salient features of transcriptional activation. The concentration of a target gene (y axis, arbitrary units) varies as a function of the concentration of an upstream transcriptional activator (x axis, arbitrary units). Three features used to summarize the dynamics of transcriptional response are highlighted: the effectiveness is reported by the orange line, the highest level of expression (HLE) is reported by the pink line, and the lowest level of expression (LLE) is reported by the green line. **(C) Net transcriptional inhibition.** The salient feature of a transcriptional inhibition is reported using the same conventions of panel B. **(D) Net response in an LRC of transcriptional inhibitors.** Responses of the genes in a LRC formed by transcriptional inhibitors are reported using conventions comparable to those of panels (B) and (C). Note i) how inhibition and activation alternate and ii) the changes in LLE (dotted blue line), HLE (dotted red line), and effectiveness (orange line).

### Simulations of LRCs and comparison to experiments on *E. coli*

The behaviour of each step of transcriptional regulation is characterised by a potentially unique combination of biological factors that control how the concentration of the source TF affects the expression of the target TF. In our model, this behaviour is controlled by four independent parameters (See Supplementary Figure 1). Our parameterization allows for greater flexibility, but limits our ability to perform a full analytical analysis. Therefore, we decided to study the behaviour of LRCs by analysing the outcome of 10000 simulations constructed by randomly sampling all the necessary parameters and measuring LLE, HLE and RE as a function of the length of the chain (up to chain length of ten). The results of our analysis are summarized in Figure 2A-C.

**Figure 2.**
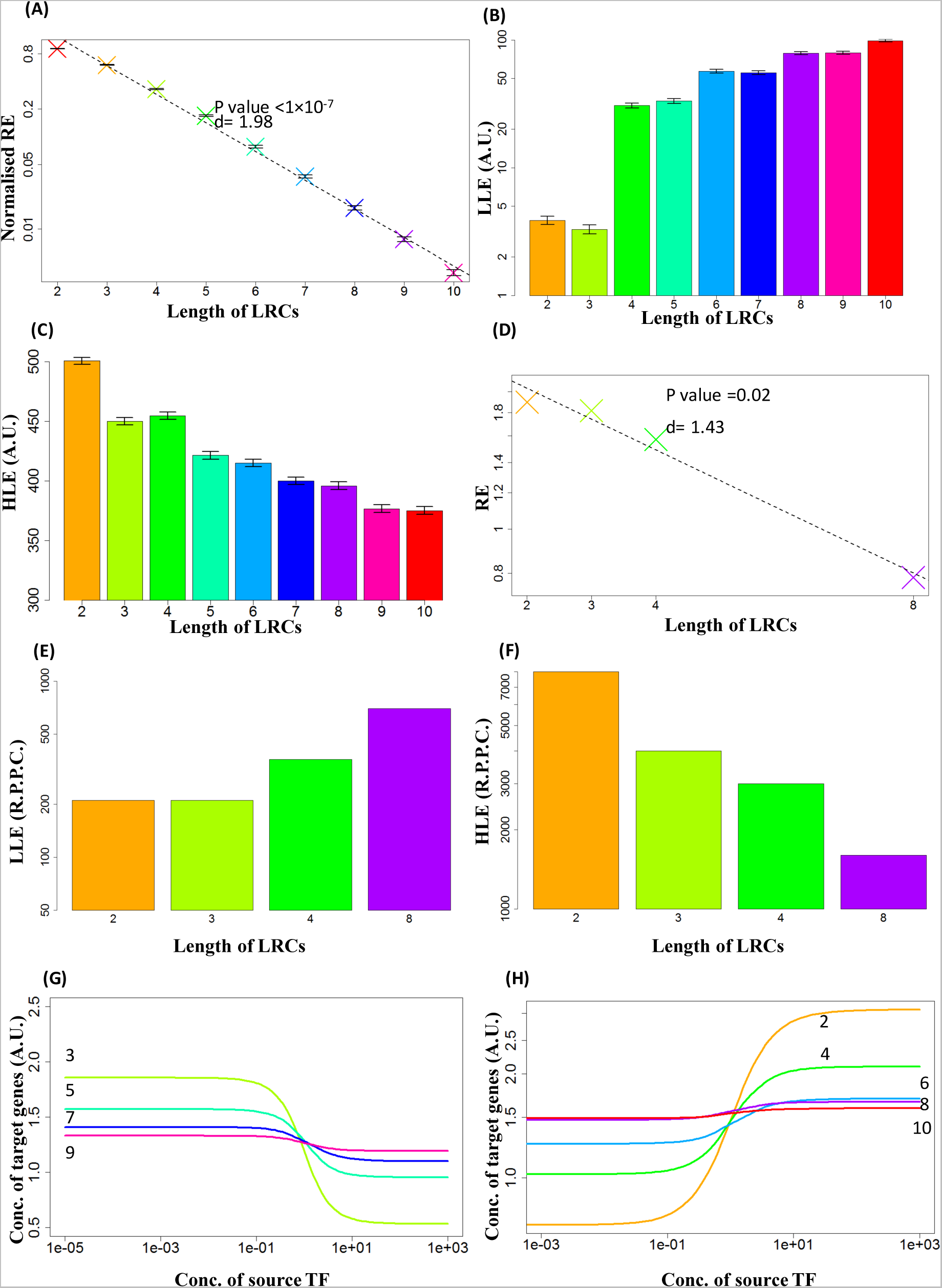
Properties of simulated LRCs and comparison with experimental data. (A) RE for LRCs of inhibitor. Mean RE across 10 000 simulations is reported, on a log scale, for LRCs of different lengths composed of transcriptional inhibitors (crosses). The dashed line indicates the linear regression on the points. Note the very good fit of the model with the data (adjusted R^2^ = 0.99). Values of REs are normalised so that RE(1)=1. Errors bars indicate the standard error. “d” indicates the strength of exponential decrease (See Materials and Methods). In this and the following panels LRC lengths are associated with different colours. **(B) LLE for LRCs of inhibitors.** Mean LLE across 10 000 simulations is reported, on a log scale, for LRCs of different lengths composed of transcriptional inhibitors. Note the steady increase. Errors bars indicate the standard error. **(C) HLE for LRCs of inhibitors.** Mean LLE across 10 000 simulations is reported for LRCs of different length composed of transcriptional inhibitors. Note the steady decrease. Errors bars indicate the standard errors. (D). **RE computed from the synthetic biology experiments of Hooshangi *et al.* 2005[1].** The y-axis reports, on a log scale, RE. Note the good fit with an exponential decrease as predicted by the mathematical model (dashed line) with a value of “d” compatible with expectations. **(E) LLE for the biological experiments referenced in the description of panel D.** The y-axis reports, on a log scale, the number of proteins per cell (R.P.P.C.). Note the steady increase as predicted by the mathematical model (Panel B). **(F) HLE for the biological experiments referenced in the description of panel D.** Note the steady decrease as predicted by the mathematical model (Panel C). **(G) Average response function for the inhibitory regulation of LRCs.** The average transcriptional response is reported for LRCs composed of an odd number of inhibitors, which results in a net inhibition. Note the decreasing Effectiveness, the increasing LLE, and the decreasing HLE as more regulatory steps separate the TF from the source gene. **(H). Average response function for the activating regulations of LRCs.** The average transcriptional response is reported for LRCs composed of an even number of inhibitors, which results in a net activation. Note the decreasing Effectiveness, the increasing LLE, and the decreasing HLE as more regulatory steps separate the TF from the source gene.

Our simulations show quite convincingly that the average RE decreases exponentially, since log(RE_n_) decreases linearly with increasing *n* (Figure 2A). This result is robust to variation in the parameter space (Figure S. 2A-F), statistically highly significant (p-value < 10^−6^) and supported by a strong goodness of fit (adjusted R^2^ > 0.9). Additionally, our simulations show that the average LLE increases (Figure 2B) and the average HLE decreases (Figure 2C) with increasing *n*. The rate of change in LLE and HLE is less dramatic than the change in RE and more sensitive to parameter choices.

Using a synthetic biology approach, Hooshangi *et al.* constructed a genetic circuit comprising a linear chain of four transcriptional inhibitors [12]. This circuit was formed by *E. coli* genes and was inserted into live bacteria. Therefore, their data are ideal to test the predictions of our model. On deriving the value of RE from their published data we find an exponential decrease as predicted by our simulations (Figure 2D). Moreoever, on deriving LLE and HLE we find a clear increase in LLE and decrease in HLE consistent with the results of our simulation (Figures 2E and 2F).

This comparison with experimental results supports the predictions of our simulations and indicates that the signal conveyed by the LRC (i.e. the effect of variation of an upstream TF on a downstream gene) gets exponentially weaker as the length of the chain increases. To better understand if this effect was limited to a chain of inhibitors, we extended our analysis to LRCs formed only by activators and to LRCs formed by a mixture of activators and inhibitors. These simulations show that our conclusions on the behaviour of RE, LLE and HLE are robust to the type of chain considered (Figure S. 1B-C).

These results indicate that in an LRC the response of a gene to the variation in the concentration of an upstream TF becomes exponentially weaker as the number of links separating them increases, and thus a long LRC acts as an *attenuator* of upstream variation. The steady increase in LLE indicates that even though each inhibitory link of a LRC may be capable of perfect inhibtion, the net inhibition of a downstream gene becomes increasingly “leaky”. Similar considerations suggest that activated genes downstream of a long LRC are only able to achieve an *imperfect activation*. These observations are supported by the behaviour of the average response function for inhibiting and activating chains (Figures 2 G-H).

This average behaviour is, however, not always observed: in a small percentage of cases (<0.5%) RE remain stable or even increase. This indicates that examples with non-decreasing RE are possible as long as the factor controlling transcriptional regulation are constrained to specific values that remains shielded from molecular noise.

### Computational analysis of LRCs in *E coli*

The results reported in the previous section support the idea that LRCs more than a few genes in length act as very strong attenuators of variation. LRCs beyond a few steps in length should be rare in real organisms since the attenuation saturates exponentially as a function of the number of links in the LRC (i.e. increasingly long chains act as increasingly imperfect regulators and exhibit increasingly lower relative effectiveness). To test this hypothesis we constructed various GRNs from the literature and computed the number of LRCs of different length. The *E. coli* GRN, obtainable from the RegulonDB database [13], is one of the most validated in the literature. Our analysis on this GRN indicates that LRCs are preferentially short and that chains with more than six genes are very uncommon (Figure 3A and Table 2). The lack of long LRCs is particularly evident when the real GRN is compared to both random (Figure 3A) and randomised (Figure S.4) networks.

**Table 2.** Statistics for the length of LRCs across different GRNs

| Organism/Cell type | Median | Maximum |
| --- | --- | --- |
| *E. coli* | 4 | 7 |
| *M. tuberculosis* | 6 | 10 |
| *S. cerevisiae* | 6 | 11 |
| Human non-cancer (GM12878) cell line | 3 | 6 |
| Human cancer (K562 leukaemia) cell line | 8 | 15 |

A consequence of our current analysis is that transcriptional regulation in LRCs is more functional for nodes closer to the source gene. Therefore, evolutionary arguments would suggest that genes deeper in the chain require additional regulatory inputs in order to exhibit functional variation of expression. To test this hypothesis, we looked at the 52 longest LRCs of *E. coli*, which comprise six TFs and one non-TF gene. In all of these LRCs, transcriptional regulation from TFs outside of the LRCs is present, indicating that *off-chain* regulation is a common, perhaps even required, feature. If we divide the genes of these LRCs into two classes depending on their distance for the source gene, a pattern emerges. The TFs constituting the upper half of the LRCs, i.e. from the second to the fourth position, are regulated on average by only a limited number of TFs, whereas the average number of regulating TFs increases significantly from the fifth position onward (Figure 3B, p-value < 2×10^−16^), consistent with our hypothesis. The relatively small number (56) of long LRCs allows us to analyse their individual structures and functionalities. These LRCs can be grouped into three categories depending on the genes contained. The first category contains the *MarRAB* operon [14], the second category contains the *Gadx-GadW* regulon [15] and the third category contains the *RcnR-RcnA* genes [16]. As reported in Table S1, the vast majority of the TFs forming these LRCs are involved in stress and antibiotic response, suggesting that such functionalities may require the tightly controlled dynamics provided by a long LRC.

The *MarRAB* operon and *Gadx-GadW* regulon form peculiar three-gene feedback loops which, due to its high level of connectivity, is quite unlikely to emergence by chance. This configuration, which we call a “chaotic motif” (Figure S. 3A), has the potential to generate highly variable gene expression profiles [9, 17]. The chaotic dynamics allow GRNs with small differences at the level of gene expression to diverge rapidly over time. Therefore we previously proposed that the chaotic motifs identified in *E. coli* could be used to promote differences in the gene expression profile across different bacteria thus generating extensive phenotypic heterogeneity in a population and promoting the emergence of antibiotic-resistant cells [9]. In figure 3C the genes forming the only two chaotic motifs observed in *E. coli* are reported in red and all of the TFs that regulate them, either directly or indirectly, are reported in blue, violet and green.

While potentially beneficial under stress conditions, a chaotic response is presumably detrimental in a stable environment. Therefore, we expect that the chaotic motifs will be tightly controlled under normal conditions. As suggested by our analysis, long LRCs could be ideal candidates to provide such tight control. Indeed, we find that both chaotic motifs are intertwined downstream with one or more of the longest LRCs of *E. coli* (Figure 3C). Statistical analysis indicates that the probability of these motifs being embedded into such long LRCs by chance is very small (p-value = 0.004).

**Figure 3.**
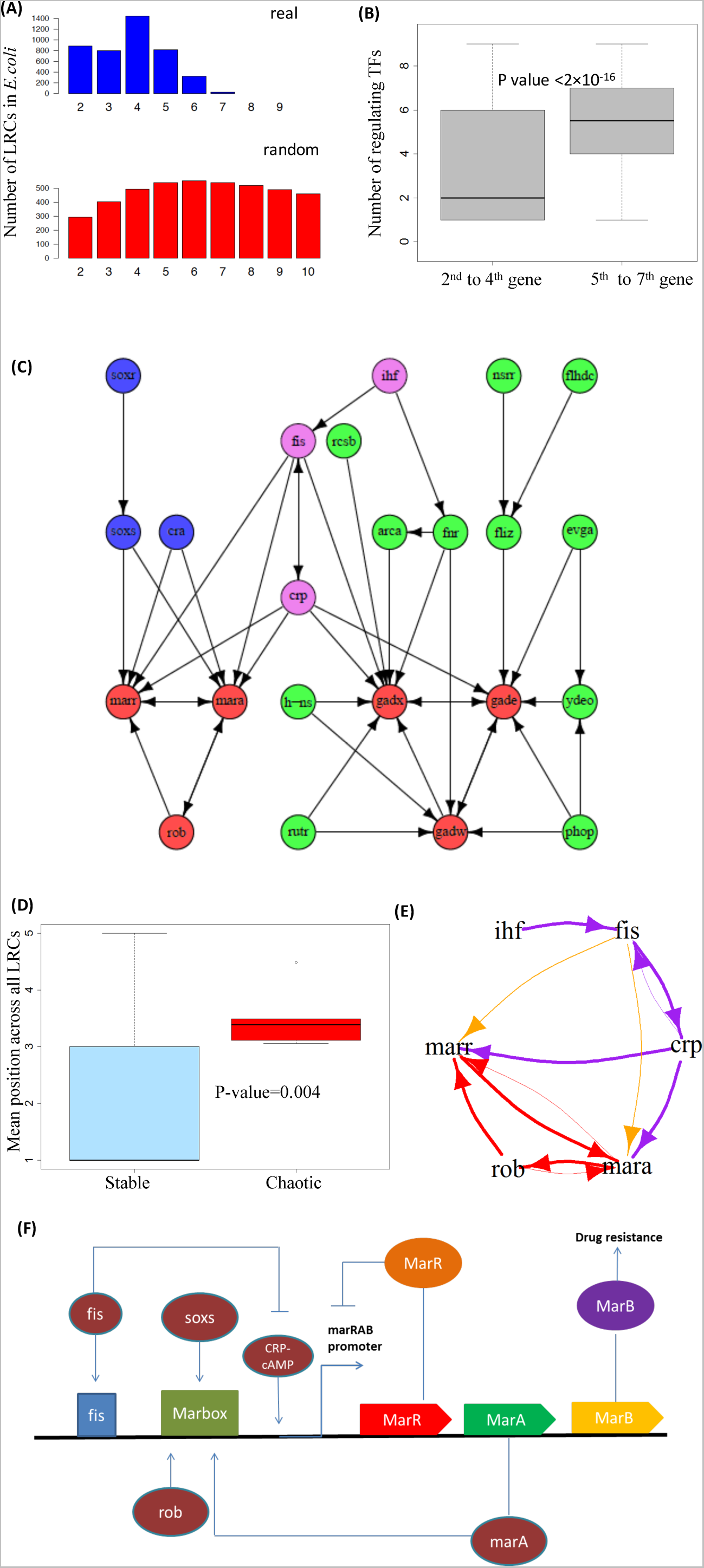
Interplay of LRCs and feedback loops in *E. coli* K12. (A) Distribution of the length of LRCs in *E. coli* K12. The number of LRCs of different lengths is reported. Note how most LRCs are formed by 5 or less genes. Length distribution of the real network is in blue and those for average random networks are in red. **(B) Degree of regulation in the longest LRCs.** The two box plots report the distribution of the number of transcriptional inputs of the genes in the upper half (2^nd^ to 4^th^) and lower half (5^th^ to 7^th^) of the longest transcriptional cascades in *E. coli*. The top genes of the chains are unregulated by construction and were not included. Note the highly statistically significant difference. (**C) Chaotic motifs in *E. coli* and their upstream regulation**. The network reports the two chaotic motifs found in *E. coli* with all the genes involved in their regulation either directly or indirectly. The genes forming the chaotic motifs are highlighted in red. The genes that control only the *Marr-Mara-Rob* motif are highlighted in blue, the genes that control only the *Gadw-Gadx-Gade* motif are highlighted in green, while the genes that regulate both are highlighted in violet. Note how the longest LRC upstream of the chaotic motifs is the shared one (violet genes). (**D) Mean position across all LRCs (MPAL) of genes involved in chaotic motifs versus other genes.** The box plots report the distribution of MPAL for each gene involved in the formation of the chaotic motifs (*Chaotic*) and for the other genes (*Stable*). Note how the MPALs of chaotic genes are significantly *larger* than the MPAL of stable genes, indicating that chaotic motif genes are encountered more frequently than average in long LRCs. **(E) Interplay of the longest LRCs and of one of the chaotic motifs of *E.coli*.** The *Marr* and *Mara* genes found in a chaotic motif (red arrows) are part of two of the longest LRCs in E. coli: *Ihf-Fis-Crp-Mara-Rob-Marr* and *Ihf-Fis-Crp-Marr-Rob-Mara* (thick violet and red arrows). Note that the motif is controlled by a 3-gene upstream LRC (*Ihf, Fis* and *Crp*) when marbox is not active (in violet), while it is regulated by a 2-gene LRC (*Ihf and Fis*) when marbox is active (see the orange arrows). A similar behaviour can be identified for the other chaotic motif. **(F) Dynamics of one of the chaotic motifs of *E.coli.*** A cartoon of the transcriptional regulation of the *marRAB* operon is plotted as derived from [2-7]. Note that *Fis* only activates the *marRAB* operon when marbox is already activated.

To further test this idea we compute the “embeddedness” of different genes into LRCs of different length. More precisely, for each gene, we computed the “mean position across all LRCs” (MPAL), which is the mean length over all the LRCs that contain that gene. Genes that are more often found in long LRCs will have a larger MPAL. Consistent with our expectation, the genes involved in chaotic motifs have a significantly larger MPAL than the other genes of the *E. coli* GRN (Figure 3D, p-value < 0.004). These results support the idea that LRC dynamics is exploited by cells to control the activation of chaotic motifs.

To explore the molecular mechanisms underpinning this theoretical prediction, we analysed the biology of the *MarRAB* operon due to the availability of extensive information on its genetics as a consequence of its key importance in antibiotic resistance [14]. The behaviour of *MarRAB* depends on the activity of the marbox enhancer DNA sequence. Experimental results indicate that *Fis* acts as a promoter of *MarRAB* only when marbox is activated by *MarA, SoxS* or *Rob* [18]. Moreover, when *MarA, SoxS* or *Rob* is absent, *Fis* reduces the activity of *MarRAB* [18]. Therefore, our theory indicates that when marbox is not active, the transcriptional activity of both *MarA* and *MarR* is tightly controlled due to the presence of a long LRC and that the potential chaotic behaviour of the motif is restricted (Figure 3E, violet arrows). Upon activation of marbox by an environmental signal – such as superoxide stress transduced through *SoxS* [19-21] – *Fis* activates the *MarRAB* operon and the LRC becomes shorter (Figures 3E, yellow arrows; Figure 3F). This allows larger variations for *MarA* and *MarB* potentially unleashing chaotic dynamics.

Comparable dynamics is observable in the other chaotic motif, which includes the genes *GadX*, *GadE*, and *GadW*. *Fis* has been reported to inhibit expression of *Gadx* in the late stages of exponential growth, when a dramatic shift in gene regulation can be observed with respect to earlier stages [22]. Interestingly, the late stages of exponential growth are commonly associated with stress [22] and the GRN of *E. coli* suggests that during this stage *Gadx* is regulated by a two-gene LRC, which is formed by *Ihf* and *Fis*, instead of the three-gene LRC which is active in the previous stages (formed by *Ihf, Fis* and *Crp*). To the best of our knowledge, the exact molecular mechanism underlying this switch has not yet been elucidated. However, our analysis provides new ways to approach the investigation of this problem [22].

### Computational analysis of LRCs in other microorganisms

The *E. coli* GRN is one of the most experimentally validated available in the literature and extensive analysis is possible. Other less well-characterised GRNs are available for *M. tuberculosis* and *S. cerevisiae*, allowing an admittedly more limited exploration. As our theory will be very sensitive to false positives, the outcome of our analysis for these organisms is potentially less robust.

Many similarities are found between the GRNs of *M. tuberculosis* and *E. coli*. The LRCs of *M. tuberculosis* are limited in number and preferentially short (Figure 4A), LRCs are shorter than expected from random networks, and the number of transcriptional regulators is significantly higher for genes deeper in the longest LRCs (Figure 4B). Remarkably, two chaotic motifs with the same structure as those encountered in *E. coli* can be found (Figure 4C Figure S. 3B), and the same intertwining of chaotic motifs and LRCs discussed above can be observed. Both of the chaotic motifs of *M. tuberculosis* are embedded into the longest LRCs of the organism (Figure 4C) and the probability of these motifs being embedded into such long LRCs by chance is small (p-value < 10^−6^). In striking similarity to *E. coli*, the two chaotic motifs in *M. tuberculosis* are controlled by LRCs comprising four genes, hinting that a four-gene LRC acts as a “universal attenuator.”

**Figure 4.**
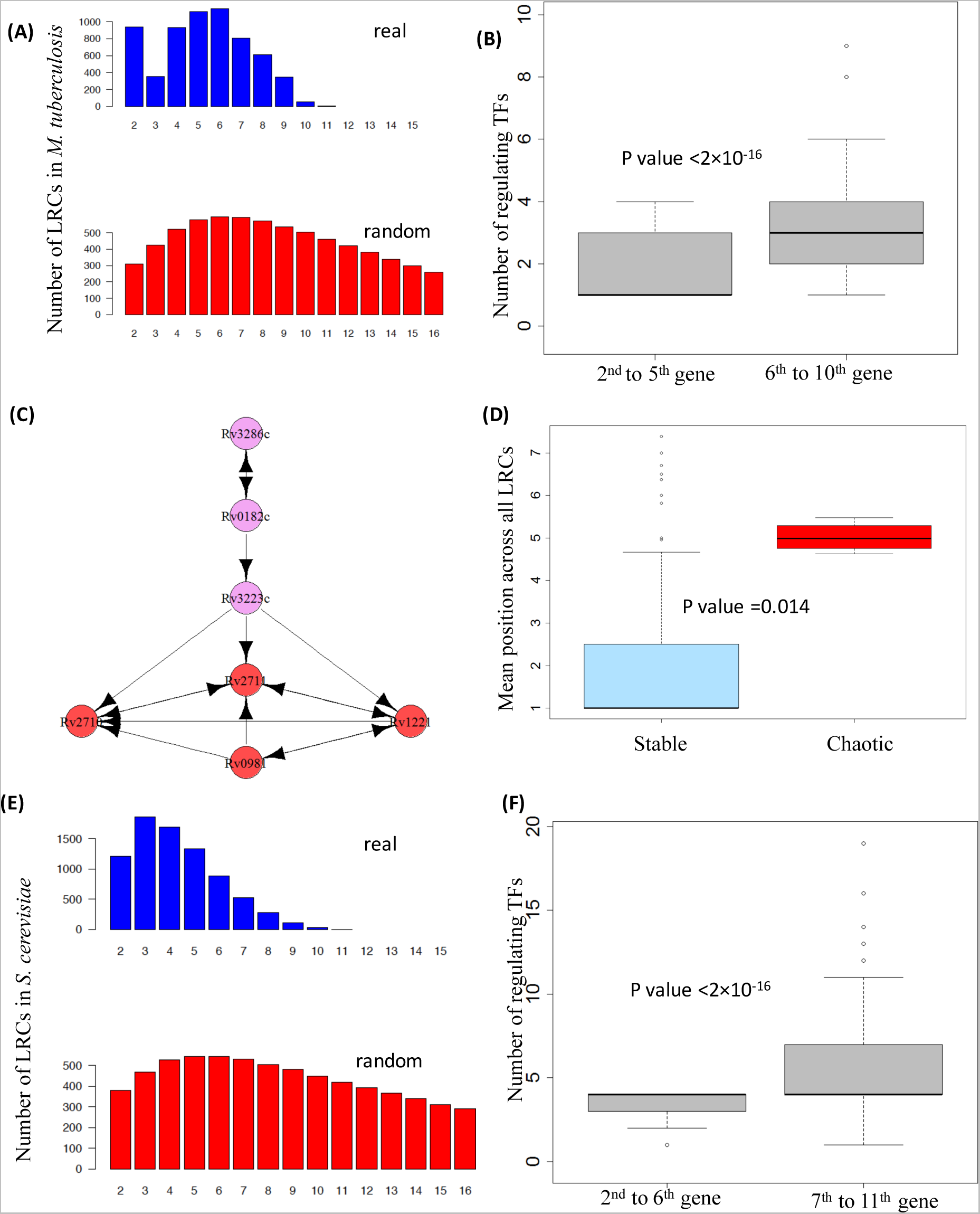
Interplay of LRCs and feedback loops in *M. tuberculosis* and *S. cerevisiae*. (A) Distribution of the length of LRCs in *M. tuberculosis*. The number of LRCs of different lengths is reported. Note how most LRCs are formed by 7 or less genes. Length distribution of the real network is in blue and those for average random networks are in red. **(B) Degree of regulation in the longest LRCs of *M. tuberculosis*.** The two box plots report the distribution of the number of transcriptional inputs of the genes in the upper half (2^nd^ to 4^th^) and lower half (5^th^ to 7^th^) of the longest transcriptional cascades in *M. tuberculosis*. The top genes of the chains are unregulated by construction and were not included. Note the highly statistically significant difference. **(C) Chaotic motifs in *M. tuberculosis* and their upstream regulation**. The network reports the two chaotic motifs found in *M. tuberculosis* with all the genes involved in their regulation either directly or indirectly. The genes forming the chaotic motifs are highlighted in red, while the genes that regulate them are highlighted in purple. Note how the LRC upstream of the chaotic motifs is the composed by three genes as in *E. coli*. (**D) Mean position across all LRCs (MPAL) of genes involved in the chaotic motif versus the other genes in *M. tuberculosis*.** The box plots report the distribution of MPAL for each gene involved in the formation of the chaotic motifs (*Chaotic*) and for the other genes (*Stable*). In agreement with expectation, chaotic motif genes show significantly higher MPAL compared with others. **(E) Distribution the length of LRCs in *S. cerevisiae***. The number of LRCs of different lengths is reported. Note how most LRCs are formed by 8 or less genes. **(F). Degree of regulation in the longest LRCs in *S. cerevisiae*.** The two box plots report the distribution of the number of transcriptional inputs of the genes in the upper half (7^th^ to 11^th^) and lower half (2^nd^ to 6^th^) of the longest transcriptional cascades in *S. cerevisiae*. The top genes of the chains are unregulated by construction and were not included. Note the highly statistically significant difference.

The experimental work of Harbison et al. provides one of the most reliable sources for transcriptional interactions in yeast [9, 23] and we reconstructed the GRN of *S. cerevisiae* from their data. However, it must be noted that other datasets exist with different properties [24, 25], thus indicating the difficulty associated with the experimental derivation of the GRN for this organism and suggesting a perceivable level of noise even in the data that we used. LRCs of moderate length are present in this organism (Figure 4E). Nonetheless, they are shorter than expected from random networks. Moreover, compatible with our expectations, the number of transcriptional regulators is higher for genes deeper in the longest LRCs (Figure 4F). No chaotic motifs are observed in this GRN.

### Computational analysis of LRCs in human cell lines

The work of the ENCODE consortium allowed the derivation of a partial GRN for two human cell lines: GM12878 and K562. The GRN of the human non-cancer cell line GM12878 behaves as expected from our theory: LRCs are preferentially short and the longest transcriptional chains consist of only four TFs and one non-TF gene (Figure 5A and C). Moreover, LRCs are shorter than expected from random networks and the number of transcriptional regulators is higher for genes deeper in the longest LRCs (Figure 5C). Compatible with the idea of a relatively stable phenotype of normal cells, no potentially chaotic motifs can be identified in this cell line.

**Figure 5.**
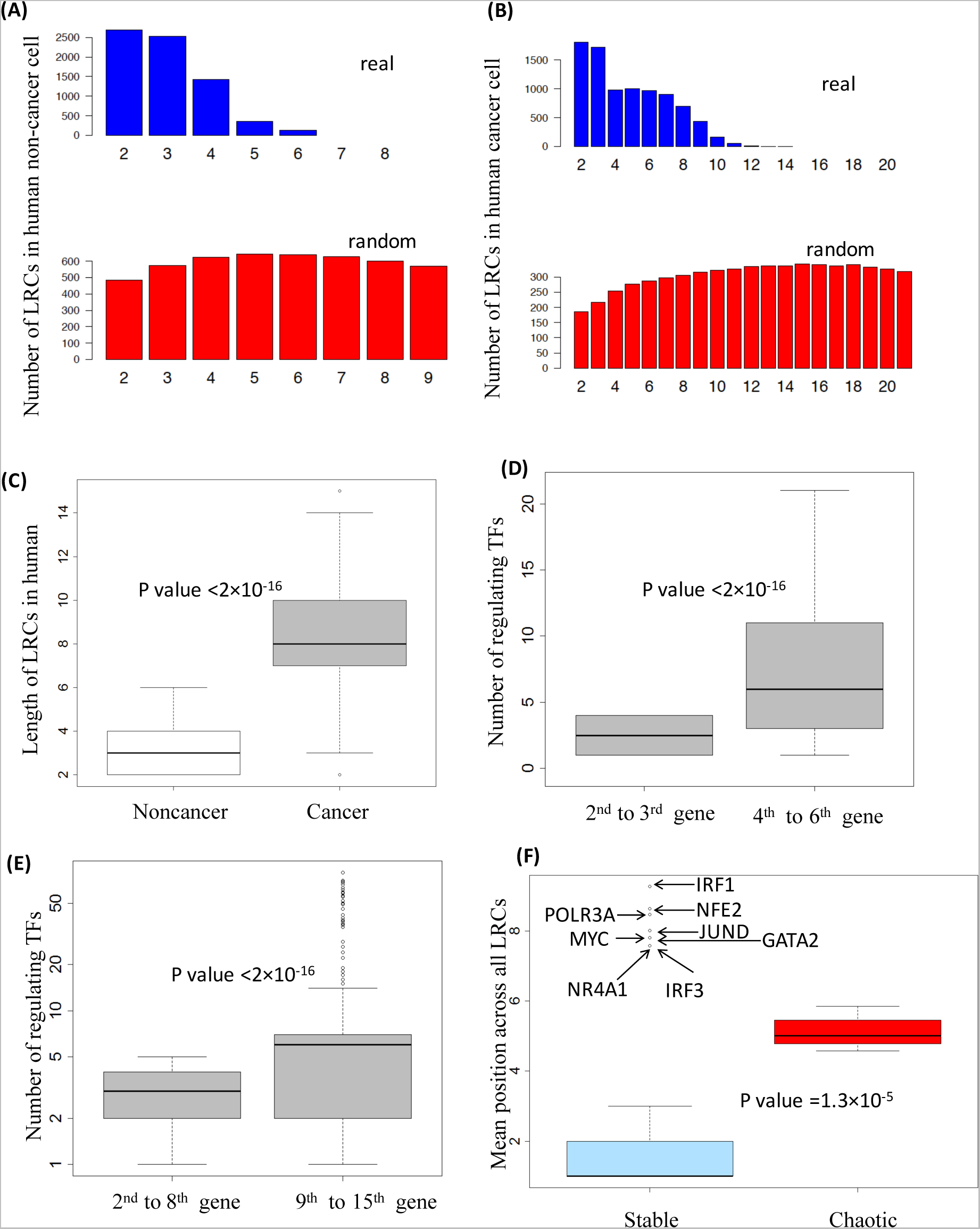
Interplay of LRCs and feedback loops in human cell lines. (A,B,C) Distribution of the length of LRCs in the two human cell lines GM12878 (non-cancer) and K562 (leukaemia). The number of LRCs of different lengths is reported. Note the striking difference in the number and length of LRCs across the different cell lines. **(D, E) Degree of regulation in the longest LRCs of human cell lines.** The two box plots report the distribution of the number of transcriptional inputs of the genes in the upper half (4^th^ to 6^th^ in GM12878 and 9^th^ to 15^th^ in K562) and lower half (2^nd^ to 3^rd^ in GM12878 and 2^nd^ to 8^th^ in K562) of the longest transcriptional cascades in the two cell lines. The top genes of the chains are unregulated by construction and were not included. Note the highly statistically significant difference. **(F) Mean position across all LRCs (MPAL) of genes involved in chaotic motifs versus the other genes in cancer**. The box plots report the distribution of MPAL for each gene involved in the formation of the chaotic motifs (*Chaotic*) and for the other genes (*Stable*). In agreement with expectation, chaotic motif genes show significantly higher MPAL compared with others. Also note how various genes implicated in cancer progression, despite not being involved in chaotic motifs, have a very high MPAL. Suggesting a narrow variation of their expression

Despite there being a comparable number of TFs and comparable link density in the two human cells lines, the GRN of the human leukaemia cell line K562 displays remarkably different properties: very long LRCs can be found, although longer LRCs would be expected from random networks (Figure 5 B, C and E), and potentially chaotic motifs consisting interlinked feedback loops are common [9, 17] (Figure 5F and Figure 6 A and B). The longest LRCs are composed of 14 TFs and one non-TF gene (Figure 5 B and C, and Figure 6 A). Interestingly, the “tail” of the longest LRCs contains several genes that are often dysregulated in cancer: *EGR1* [26, 27] [28]*, IRF3* [29]*, POLR3A* [30], and *IRF1* [31, 32].

All but one gene (*GTF2F1*) involved in the formation of potentially unstable long feedback loops are embedded into the longest LRCs found (Figure 6A). The probability of observing this embedding by chance is small (p-value < 10^−2^) and all the TFs involved in the formation of potentially chaotic feedback loops display a large MPAL (Figure 5F).

The longest LRCs have a peculiar structure. Two four-gene LRCs control the complex set of feedback loops. Remarkably, a single four-gene LRC, composed by *EGR1*, *IRF1, POLR3A* and *IRF1*, can be found in the tail of the longest LRCs (Figure 6A), emanating from the feedback loops. Our theoretical analysis suggests that the dynamics of the final gene of this chain (*IRF1*) is highly constrained, and biological experimentation indicates that *IRF1* is a tumour suppressor gene relevant to a number of cancers including leukaemia [32-34]. Our theoretical interpretation is that the dynamics of LRCs is exploited by cancer cells to inhibit the proper activity of this gene.

**Figure 6.**
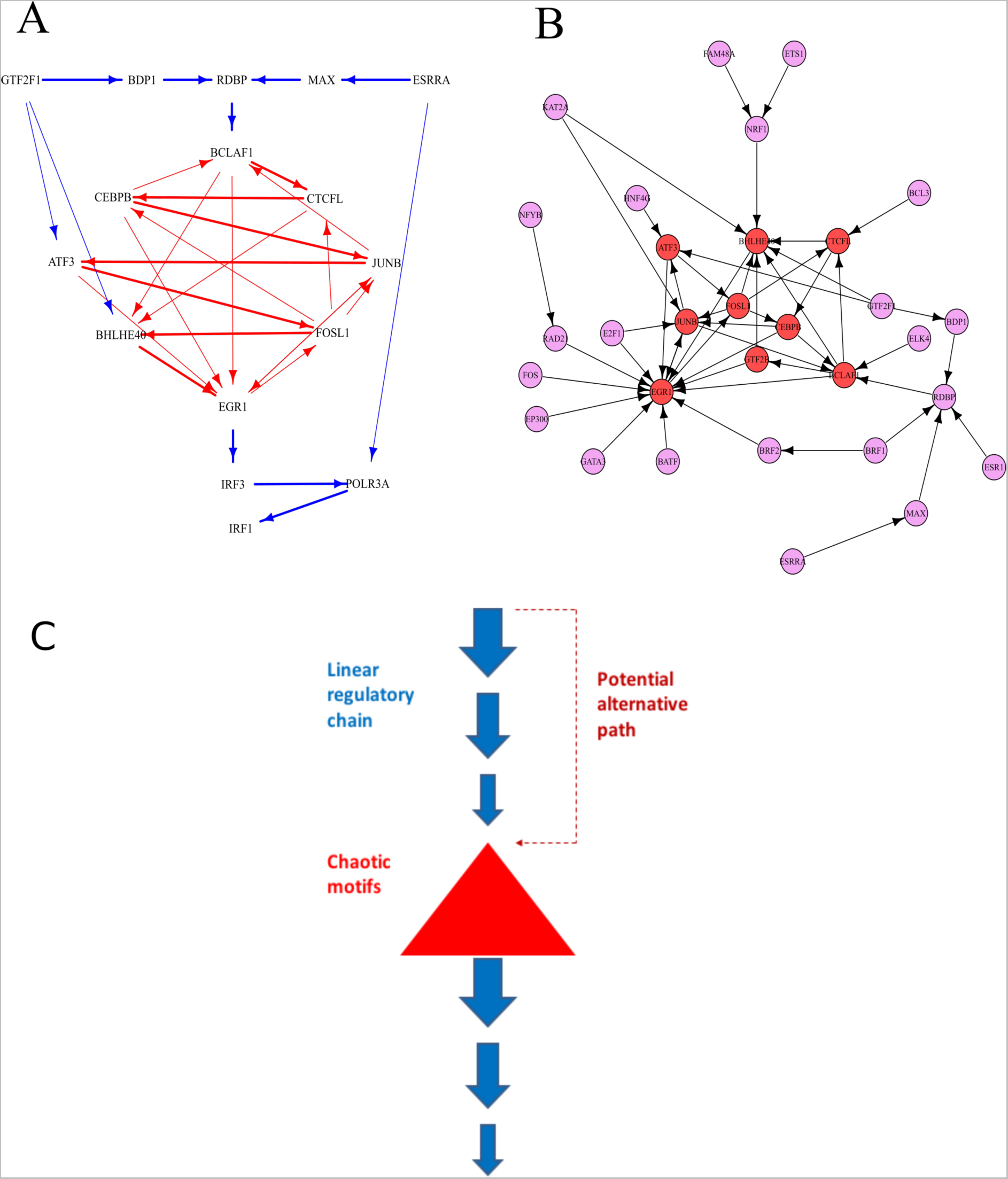
Chaotic motifs in cancer cells (K562) and their upstream regulations (A) Interplay of the longest LRCs and chaotic motifs in cancer. The network formed by all the TFs implicated in the longest LRCs of the cancer cell line K562 is reported. Note how the longest chains (thick arrows) are entangled with the feedback loops. Two 3-gene LRCs can be observed upstream of the feedback loops and one 3-gene LRC can be observed downstream (blue arrows). The presence of the 3-gene LRCs upstream of potentially chaotic motifs is consistent with our findings in *E. coli* and *M. tuberculosis*. Also note how *IRF1*, a gene which is a key regulator of growth of leukaemia and other cancer cells [8-10], is found Sdownstream of the long feedback loops. **(B) Chaotic motifs in K562 and their upstream regulation.** The network reports the genes involved in the formation of the chaotic motifs of K562 (red) with all the genes involved in their regulation either directly or indirectly (violet). Note how the chaotic motifs sit at the bottom of multiple LRCs. **(C) A cartoon demonstration of proposed functional interaction between long linear regulatory chains (LRCs) and chaotic motif.** In gene regulatory networks, upstream long LRC restricts transcriptional dynamics of chaotic motif, which has the potential to produce diverse transcriptional profiles under similar biological conditions. This restriction can be walked around via alternative regulatory path in certain organism such as *E. coli* upon activation. In addition, Long LRC can be used to insulate important downstream gene from the dynamics of chaotic motifs.

Note that a group of genes with a high MPAL can be identified in the *Stable* group. These genes are therefore likely to be encountered in relatively long LRCs, and hence conceivably have very limited RE. Notably, all of these genes have been shown to have an important role in the survival of leukaemia cells (*NFE2*[35], *POLR3A*[36], *JunD* [37], *Myc*[38], *GATA2*[39], *NR4A1*[40], *IRF3*[41], *IRF1*[32-34]) suggesting that cancer cells may be dynamically controlling the variation of these genes.

Potential biases introduced by under sampling and errors in the human GRNs limit the power of a direct mathematical approach. However, the strong diversity observed in different topological features of the non-cancer and cancer cell lines is an indication of a profound difference in their transcriptional programs. Therefore, a direct comparison between the GRNs of the two cell lines can be very informative in highlighting differences that may then be used to develop new therapies [9].

## Discussion

Theoretical and experimental efforts have provided strong evidence that variability in gene expression plays a significant role in controlling cellular phenotypes in development [42], health [43] and disease [44, 45]. While the transcriptional mechanisms responsible for controlling this variability continue to be an active area of research [46], the potential system-level interactions are less well explored, and network analysis of GRNs is a powerful approach to fill this gap. GRNs provide a description of the coordination of gene expression at a systems level and can therefore be used to explore the potential role of *topological* features that control the variability of genes expression.

### Long LRCs “pin down” relative effectiveness

Our mathematical model suggests that long LRCs “pin down” the expression of downstream genes, limiting their ability to vary in response to environmental or intracellular cues affecting the gene at the top of the chain. In fact, the variation is predicted to decay exponentially along the chain. This conclusion is supported by data derived from synthetic biology experiments on *E. coli* [12].

A direct consequence of our model is that long LRCs are ineffective in transmitting variation in gene expression beyond a few transcriptional steps. Therefore, over evolutionary time, one might argue that long LRCs yield inefficient information transmission and will have been negatively selected, resulting in relatively small numbers of long LRCs in GRNs. This prediction is supported by an analysis of the GRNs of different organisms, ranging from bacteria (*E. coli* and *M. tuberculosis*), to yeast and human.

It has been observed that the sensitivity of gene regulation becomes higher as LRCs get longer and a “switch-like” behaviour is observed; this can be interpreted as the result of an increasingly larger effective Hill-coefficient [12]. Whilst our findings do not contradict this, we present an additional observation that the terminal gene of a long LRC will display only a limited range of variation in response to changes in the concentration of the source gene. Beyond a length of approximately four links, due to the exponential decay, the range of variation is very small and likely to be comparable in magnitute to the fluctuations in gene expression due to intrinsic molecular noise. In addition, our modelling indicates that the effectiveness of regulation is compromised by the LRC topology itself. For example, a linear chain of an odd number of perfect inhibitors will have a net effect of imperfect inhibition, and the degree of imperfection will increase with the length of the chain. Taken together, these observations imply a tradeoff between the sharpness and the effectiveness of net regulation through a LRC, which depends on the specific parameters that characterize the interactions, but nonetheless strongly suggest that very long LRCs are of limited utility in GRNs, and hence negatively selected through evolution.

Our model also suggests that the average behaviour of LRCs can be prevented by constraining the biological parameters associated with transcription to very specific values. This suggests that, under specific circumstances, biological processes may be in place to prevent the emergence of such average behaviour in LRC. The precise regulation of the parameters required also suggests that molecular insults are very likely to push LRCs towards the average expectation, potentially changing the behaviour of cells.

### Chaotic motifs are potential drivers for heterogeneity

Under normal conditions, cells must be able to filter the fluctuations of protein concentration, which are due to molecular noise. To this end, they need to display a stable response. A consequence of this type of response is the limitation of the heterogeneity of a population of cells, as each cell exposed to similar stimuli will react in a comparable way. Therefore, the very same stable behaviour that helps cells withstand a noisy environment can be detrimental under stress condition, such as an antibiotic treatment, as in this circumstance heterogeneity is helpful in allowing the emergence of resistant subpopulations of cells.

Therefore, it has been suggested that network motifs in the GRNs of bacteria can be activated only when the cell is exposed to stress [19-21]. Ideally, these motifs should have the potential to produce chaos. Chaos theory is a well-known mathematical theory that studies the behaviour of systems that are extremely sensitive to initial conditions — a paradigm popularized by the so-called “butterfly effect”. In a chaotic system, small differences in initial conditions can yield widely diverging states after a relatively short time [47, 48].

Theoretical studies indicate that certain network motifs have the potential to produce a chaotic response [17] and recent experimental work has shown complex oscillations and, loosely speaking, chaotic dynamics of certain GRN motifs both in cell-free system and *in vivo* [49]. Since a chaotic response is able to generate wildly different values by starting from very similar initial conditions, it has been suggested that chaos can act as a “heterogeneity engine” that allows a population of cells to quickly explore a large number of phenotypes [9]. Such phenotypic heterogeneity is likely to play a crucial role in allowing the emergence of resistant clones which will help a population to overcome challenging conditions such as environmental stress and antibiotic treatments [50, 51].

As discussed above, minimal chaotic motifs can be identified in the GRNs of *E. coli* and *M. tuberculosis*. Moreover, more complex and somewhat more disorganized chaotic motifs can be found in cancer. This suggests a strong parallelism between the systemic processes that allow bacteria and cancer to generate heterogeneity and ultimately to overcome the ability of the immune systems to properly fight infections and cancer.

### Long LRCs suppress generators of potential “butterfly effects”

A limited number of long LRCs can be observed in the GRNs analysed. This suggests that such configurations may be important to limit the gene expression level of few selected genes. Remarkably, we found that in both *E. coli* and *M. tuberculosis*, long LRCs are associated with genes activated during stress and antibiotic response. The expression of stress response genes is associated with an increased metabolic cost, which generally results in a reduced growth rate [52-54]. Therefore, it is reasonable to expect a tight control of these genes to prevent a dampening of the fitness of a population. Indeed, such tight control is embodied in the dynamics of LRCs. Additionally, the transcriptional control exerted by LRCs on genes downstream in the chain can help reduce noise arising from stochastic gene expression and fluctuations in the cellular environment [55-58].

The entanglement of long LRCs with potentially chaotic motifs in *E. coli* and *M. tuberculosis* suggests that the dynamics of long LRCs may allow these organisms to directly influence phenotypic variability and hence population-level heterogeneity by allowing a chaotic response only when needed. This finding is supported by the biology of both the *MarRAB* operon and the *GadW*, *GadX,* and *GadE* genes in *E. coli* and suggests new ways to bolster the effectiveness of drug treatments by targeting the mechanisms that lead to the emergence of resistant clones in bacterial populations. *In all the GRNs that we analysed, the chaotic motifs are observed after LRCs composed of exactly four genes.* This is in remarkable agreement with our theoretical observation that the genetic variation is tightly restrained from the fourth gene onward, and leads us to propose that four-gene LRCs act as “universal attenuators”.

Long LRCs and potential chaotic motifs are entangled in such a way to support both strong variations in the expression of certain genes, i.e. those within long feedback loops, such as *EGR1* (a regulator of multiple tumour suppressor genes [59]) and a very limited variation in the expression of others, i.e. those residing at the end of LRCs, such as *IRF1* (an essential regulator of growth of leukaemia and other cancer cell types [31-33]). Indeed, four-gene LRCs operate at both the “input” and “output” of non-linear feedback loops in the K562 GRN. The combined action of these competing dynamics may be able to generate heterogeneity while limiting the necessary variation in gene expression associated with tumour suppression.

Our findings may provide a mechanistic basis for “oncogene addiction” [60, 61]. The term is used to indicate that some tumours depend on the constitutive activation of a single oncogene for sustaining growth and proliferation and that transient inactivation of that particular oncogene may be enough to promote differentiation or apoptosis of cancer cells [62]. Universal attenuators may drive the constitutive activation of a gene, and thus targeting of LRCs could be a novel strategy for cancer cell killing.

### Analysis of LRCs shed new light on the topological pressure acting on GRNs

We have previously shown that mathematical modelling can be used to explore the topological features associated with robustness in GRNs. In particular, the theory of Buffered Qualitative Stability (BQS) postulates that long causal chains of genes, irrespective of the in-degree of the gene at the top of the chains, should be limited in number due to their evolutionary susceptibility to seeding long feedback loops, which can create instability[9]. Taken together with our current result, this indicates that long causal chains of TFs are dangerous and with limited functionalities, thus suggesting that healthy cells should have very limited instances of such configurations. This is indeed observed in real data.

Further connections emerge when potential sources of instability (chaotic motifs) are contextualised with respect to LRCs. When chaotic motifs are identified in a GRN, they are entangled downstream of long LRCs. Moreover, and quite unexpectedly, an LRC comprising exactly four genes (and therefore three transcriptional interactions) can be found upstream of all the chaotic motifs. This strongly suggests that four-gene LRC provides a general mechanism in GRNs to ‘pin down’ or insulate the genes involved in the generation of a chaotic response, hence allowing a topological control of heterogeneity.

## Conclusions

We have presented a set of results arising from theoretical modelling, statistical simulations and data analysis, all focused on the role of two different topologies in GRNs, namely, long linear regulatory chains (LRCs) and chaotic motifs. Our modelling work indicates that LRCs have a key role in reducing variation in gene expression, while chaotic motifs can act in the opposite manner and generate strong variation through chaotic dynamics. LRCs are highly effective at shutting down variation, and hence there is no additional benefit for a GRN to have very long chains, a result which is consistent with the GRNs analysed. Chaotic motifs, in being able to generate variation so rapidly, would presumably be inactivated in the steady state of a cell’s life cycle, and indeed we find in bacteria and a human non-cancer cell line that such motifs, when present, always sit at the end of relatively long LRCs, implying that they are strongly suppressed. The GRN of a human cancer cell line exhibits a much richer interplay between LRCs and chaotic motifs, and we postulate this may allow a given cancer cell to drive strong variation in certain genes and inhibit expression of tumour suppression genes, thereby allowing optimal conditions for growth and survival in the challenging environment of host tissue. Due to the ubiquity in the GRNs studied of four-gene LRCs, we postulate these modules as “universal attenuators”, with a key role of controlling potentially chaotic feedback loops.

Our work provides evidence that one can exploit knowledge of the topology of GRNs to exert a direct control on the variability of genes, even if a precise characterization of the parameters that control gene regulation is unavailable. Given the qualitative differences between the GRN topologies of normal and cancer cells [9], this may provide a way to design new targeted therapies that selectively affect gene expression variability only in cancer cells.

## Materials and Methods

### Mathematical model of linear regulatory chains (LRCs)

Each gene forming a given LRC was associated with a value that identifies both its expression level and the concentration of its transcribed protein. Moreover, we assumed the delays due to transcription initiation and translation to be negligible. Additionally, we assumed the linear chain to be autonomous, i.e. the concentration of a gene depends directly only on the concentration of the gene directly upstream. Therefore, the level of expression of each gene in the chain depends, directly or indirectly, on the concentration of the gene at the top of the LRC. The interaction among sequential genes was modelled by a deterministic interaction function. In particular, by assuming steady state dynamics, we can describe the concentration of gene *y* in the LRC as

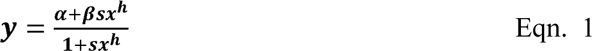

In this equation, *x* is the concentration of the transcriptional regulator directly upstream of the gene under consideration. The parameters of the equation characterize the mode and intensity of the interaction: a models the concentration of *y* when *x* is not present, /3 models the concentration of *y* when *x* is highly expressed, *h* is the Hill coefficient which describes the “cooperativity” of transcriptional regulation, and *s* is a quantity associated with the shape of the interaction function. When β>α, the equation describes a transcriptional activator and when β<α, it describes a transcriptional inhibitor (Figure S.1A and Figure S. 1 D-G).

### Quantitation of general properties of transcriptional regulation

The interaction term described by Eqn. 1 was used to model the response of the terminal gene of a LRC with a different number of genes when the concentration of the gene at the top of the LRC is varied between zero and infinity. To this end, we introduced three measures relative to the response function of the terminal gene: “Relative Effectiveness” (RE), “Lowest Level of Expression” (LLE), and “Highest Level of Expression” (HLE). These quantities are described in the main text and in Table 1.

### Simulation of LRCs and sensitivity analysis

The RE, LLE and HLE of 10000 simulated LRCs with 2 to 10 genes were computed. Both concentration and parameter values were measured in arbitrary units. For each gene of the LRC, the parameters that control the interactions were randomly generated using a uniform sampling. The parameters α, β were sampled between 0 and 1000 to indicate up to a 1000-fold activation or inhibition, *h* was sampled between 1 and 10 to account for polymeric regulation of up to 10 transctiption factors and *s* was sampled between 0 and 10 to account for different activation thresholds. For all the sampling ranges, the boundaries were excluded. Simulations and statistical analyses were performed in R version 3.2.2. To assess the sensitivity of our results to different parameters values, we performed an exhaustive computational analysis to explore the outcome of our analysis when different ranges for the parameter were used (Figure S. 2). This analysis supports the robustness of our conclusions.

### Assessment of the exponential decrease of RE

To formally assess the exponential decrease of RE, we used a logarithmic transformation. If a quantity decreases exponentially, at each step the previous value is divided by a constant (*d*). Therefore, calling RE_n_ the average RE after *n* regulatory interactions, we have:

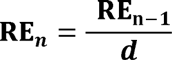

And therefore

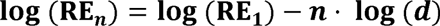

We used a linear regression on the log-transformed data and computed the p-value for the estimation of the slope and the adjusted R^2^ for the liner model. A p-value close to zero indicates that the slope is significantly different from zero and therefore that a clear exponential decay was present in the non-log-transformed data. An adjusted R^2^ close to 1 indicate that the lineal model describes the data very well. Therefore, the values reported in the main text support the existence of a strong exponential trend.

### Graph manipulation and analysis

GRNs described in the main text were derived with the same procedures and parameters described in a previous work [9]. In particular, the E. coli GNR was obtained from RegulonDB version 8 [13] by considering only interactions supported by at least two evidence codes, the *M. tuberculosis* GRN was obtained from literature [63] by considering all interactions, the yeast GRN was constructed from literature by considering interactions obtained under rich media growth supported by a p-value lower than 10^−3^, the human non-cancer and cancer cell GRNs were constructed from the proximal filtered network derived from the ENCODE data [64] for the GM12878 and K562 cell lines respectively. See our previous work [9] for a discussion on the rationale behind these choices.

The number of LRCs can be very high in complex networks with a high edge density. Therefore, we decided to develop a probabilistic algorithm to sample from the complete set of LRCs. Specifically, an LRC was grown from a starting node *S* selected with a probability proportional to its total degree. Starting from S a biased random walk was performed by randomly selecting an upstream node (with a probability proportional to its in-degree) or a downstream node (with a probability proportional to its out-degree). Nodes already present in the LRC were excluded from further sampling. This growing procedure was repeated until no upstream or downstream nodes were available. 10000 LRC samples were considered for each network and duplicates were removed from the count. The procedure described along with all the other operations on the networks were implemented using R version 3.2.2 and the “igraph” package version 1.0.1 [65]. The code is available as a supplementary file.

To perform the statistical analysis of the embeddedness of chaotic motifs with the aim of assessing the probability of interaction of the chaotic motifs with the longest LRCs of the network we employed a simple statistical model that could avoid many of the mathematical and computational complications arising from the comparison of real networks with a randomised null model. In particular, we focused on the behaviour of the TF directly upstream of the chaotic motifs, which will be referred as *U*. In all of the GRNs considered in this article, this *U* is the third gene of the longest LRCs. *U* regulates n other TFs. Of these, n_c_are chaotic TFs, i.e. TFs that take part in the formation of chaotic motifs, and n_s_ =n - n_c_ are stable TFs, i.e. TFs that *do not* take part in the formation of chaotic motifs. Due to the way in which GRNs have been constructed, *U* cannot regulate itself. Moreover, due to the structure of LRCs *U* cannot regulate the two TFs upstream in the longest LRCs. It is important to stress that these two upstream TFs are not chaotic. Therefore, if a GRN contains N_TF_ TFs, U could in principle regulate up to N_TF_ -3 TFs. The N_TF_ genes can be further divided into N_C_ chaotic TFs and N_S_ = N_TF_ - N_C_ stable TFs. Using a hypergeometric distribution, it is possible to compute the probability that when n TFs are selected from a set of containing N_C_ chaotic genes and N_S_ -3 = N_TF_ - N_c_ -3 stable genes at least n_c_ chaotic genes are selected. This probability represents the p-value included in the main text.

### Random and randomized GRNs

A random network associated with a GRN formed by *g* genes, *n* TFs and *e* edges was obtained by randomly placing *e* edges on an empty network with *g* nodes, in such a way that the source of each edge was randomly selected from a fixed set of *n* nodes. The randomised (rewired) GRNs were derived from original GRNs of the corresponding organisms using the “rewire.edge” function of the igraph package version 1.0.1 in R version 3.24. The number of rewiring iterations for each GRNs was set to ten times the number of edges in the network. 100 randomised GRNs were generated for each organism or cell type.

### Sources for biological data used

The values of RE, HLE and LLE along the linear transcriptional chains in the bacterium *E. coli* K12 were obtained from the experiments conducted by Hooshangi *et al.* 2005 [12]. Gene and transcription regulation data for *E. coli* K12 were obtained from RegulonDB [13] with the same procedures and parameters described before [9]. Data on the functions of genes in *E. coli* K12 were obtained from the referenced literature and the EcoCyc web resource [66, 67]. The GRN for *M. tuberculosis, S. cerevisiae* and human were derived from experimental data [23, 63, 64] using the same procedures and parameters described before [9].

### Author contributions

DL drafted the manuscript, provided original concepts, designed the computational experiments, designed and implemented the simulations and performed the statistical analysis. LA drafted the manuscript, provided original concepts, designed and implemented the computational experiments and performed the statistical analysis. TN drafted the manuscript, provided original concepts and performed analytic calculations.

## Acknowledgements

The authors thank Md. Al Mamun and Julian Blow for helpful comments throughout the project. DL acknowledges support from the Wellcome Trust PhD programme. LA and TJN acknowledge partial support from the Scottish University Life Science Alliance. The authors also acknowledge High Performance Computer resources partially supported by the Wellcome Trust (Centre Grant 083524).

## Footnotes

1 Transcription factors that are only regulated by feedback loops are also considered at top layer. In the GRNs analysed in this study, there are very few transcription factors in this category.

## List of Abbreviations

TF: Transcription factor
GRN: Gene regulatory network
LRC: Linear regulatory chain
RE: Relative effectiveness
LLE: Lowest level of expression
HLE: Highest level of expression
MPAL: Mean position across all LRCs

